# Pan-ASLM: a high-resolution and large field-of-view light sheet microscope for Expansion Microscopy

**DOI:** 10.1101/2025.08.06.668765

**Authors:** Hannahmariam T. Mekbib, Lasse Pærgård Andersen, Shuwen Zhang, Jonathan Gulcicek, Yuan Tian, Jack R. Ross, Mark D. Lessard, Joerg Bewersdorf

## Abstract

Expansion microscopy, a super-resolution fluorescence microscopy technique in which samples are expanded up to ∼8,000 times (after 20-fold expansion) their original volume, places high demands on the microscopes used to image the expanded samples. To reveal nanoscale cellular ultrastructure in meaningful sample volumes, the instruments need to feature a large field of view and working distance. Simultaneously, they need to offer a high three-dimensional resolution to avoid counteracting the resolution improvement achieved by the expansion process. Here, we present pan-ASLM, a high resolution, large field-of-view light-sheet microscope developed for expanded samples, based on the Axially Swept Light Sheet Microscopy (ASLM) technique. pan-ASLM allows imaging over a 640 µm x 640 µm field of view with lateral and axial resolutions of 566 nm and 457 nm, respectively, and features an image acquisition speed of up to 20 fps (183 Mvoxels/sec). It offers ∼1200x higher imaging speed, a ∼7x larger field of view, and ∼2x better axial resolution than the standard confocal microscopes typically used for expanded samples. We validate the new microscope design through imaging of pan-expanded HeLa cells as well as mouse kidney and brain tissue.

## Introduction

Fluorescence microscopy [1] is the imaging technology of choice for how biomedical researchers investigate the distribution of molecules of interest in cells and tissues. Super-resolution microscopy techniques developed over the last few decades have overcome the diffraction limit of resolution of ∼250 nm, achieving resolutions of down to ∼20 nm and better [2–5]. Among these techniques, Expansion Microscopy (ExM) [6–11] differs by its means of resolution improvement: rather than improving the resolution by utilizing optics and/or the switching properties of fluorescent labels, ExM physically expands biological samples by embedding them in a hydrogel that physically swells by a factor of ∼4 to ∼20 in each dimension. The embedded structures grow in size by the same factor, which leads to an equivalent effective resolution improvement. We have recently developed pan-Expansion Microscopy (pan-ExM) [12–14], a variant of ExM which provides ∼13-24x expansion while retaining proteins in bulk. Labeling the retained proteins with a fluorescent dye reveals the cellular ultrastructure, providing images resembling those obtained by Correlative Light and Electron Microscopy (CLEM) but using just a conventional light microscope. The large expansion factor of pan-ExM and other ExM methods amplifies, however, a major challenge many modern microscopy applications face: increasingly, researchers seek to image whole tissue sections in three dimensions (3D) at high spatial resolution and sufficiently high throughput. This simultaneous need for a large field of view (FOV), long working distance (WD) of the objective lens, high resolution, and fast imaging speed is much harder to meet when the sample has increased 8,000-fold (20-fold linearly expanded) in volume.

Conventionally, point-scanning confocal microscopes [15] have been used to image expanded samples as they provide high-contrast images by using a pinhole that prevents out-of-focus light from reaching the photodetector [16]. The achievable resolution of these instruments is ∼250 nm in the focal plane (XY) and ∼800 nm axially (Z) when using high-NA water immersion objectives (e.g. 60x/1.2 NA). This translates to ∼13 nm XY and ∼40 nm Z resolution after correcting for a 20x expansion factor. However, for larger FOVs (e.g. 200×200 µm^2^, which still only corresponds to a sample area of 10×10 µm^2^ before expansion), acquiring a 3D stack becomes a serious bottleneck for this imaging technique: because of the low signal level in expanded samples (less molecules per unit volume due to the expansion) extensive averaging is usually required. This leads to very long imaging times of tens of seconds for a single Nyquist-sampling limited large-FOV image and results in hours of acquisition time for a single 3D stack of a few hundred images.

Spinning disk confocal microscopes [17] can speed up the imaging time substantially by parallelizing the scan process using many pinholes simultaneously. However, diffraction-limited performance with these microscopes is only achieved with high-magnification, high-NA objective lenses due to optical constraints in the microlens disk design which limits high-quality imaging with these instruments to small FOVs and short WDs. Low-magnification (i.e. large FOV) objectives that simultaneously feature large NAs, which would be preferred for imaging large volumes at high resolution, feature pupil diameters D so large (D=2*NA*f_TL_/M; where M is the magnification of the objective and f_TL_ the nominal focal length of the tube lens the objective is designed for) that the microlenses substantially underfill them. This compromises the resolution - for a two-fold underfilled pupil, the lateral resolution worsens by a factor of two, the axial resolution by a factor of four - and leads to the microscope further deviating from the ideal of an isotropic resolution.

Light Sheet Fluorescence Microscopy (LSFM) [18–20] presents an attractive alternative that overcomes this problem. Here the excitation and detection beam paths are separated, usually by using two objectives in an orthogonal arrangement. Creating a sheet of light for fluorescence excitation by the first objective and imaging the emitted fluorescence with the other one decouples the WD and FOV of the detection objective from the axial resolution if the light-sheet thickness is less than the depth of focus of the detection objective. Using fast cameras, LSFMs can therefore theoretically image large FOVs at high speeds and axial resolutions exceeding those obtained by point-scanning or spinning disk confocal microscopes using the same detection objective. However, for classical (i.e. Gaussian) light sheets, there is a tradeoff between the thickness of the sheet and the axial range within which such thickness can be maintained: the thinner the sheet, the shorter the range. Since this range represents the lateral extent of the sheet from the standpoint of the detection objective, good axial resolution comes at the cost of a small FOV in which this axial resolution is achieved. To overcome this tradeoff, efforts have gone towards engineering uniformly thin light sheets with a large lateral extent [21–23].

Among those, Axially Swept Light Sheet Microscopy (ASLM) [24–26] is a particularly attractive solution. ASLM uses aberration-free remote focusing [27–28] to create uniformly thin light sheets over large FOVs. Several ASLM microscopes have been developed for specific applications over the past years, offering different combinations of resolution, FOV, and image acquisition speed [29–31]. However, none of the current ASLM designs has been optimized for pan-ExM samples, which have particularly high demands on maximizing the FOV (due to the dramatically increased sample size) and axial resolution (to not counteract the resolution improvement achieved by the expansion factor): while Glaser et al. [32] reported a very large FOV of 10.6 mm x 8 mm, the axial resolution of 3 µm has been relatively poor; SIFT [33] images an 870 µm x 870 µm FOV at the high speed of 40 fps but the isotropic resolution of 970 nm is substantially worse than what is achievable by confocal microscopes; Chakaborty et al. [25] reported ∼480 nm isotropic resolution albeit at the cost of a much smaller FOV of 320 µm x 320 µm; Lin et al. [34] reported a larger FOV ASLM variant (774 µm × 435 µm) with ∼500 nm (averaged over the FOV) resolution, high speed applications were, however, not demonstrated.

Here, we introduce pan-ASLM, a new LSFM specifically developed and optimized for high-throughput imaging of highly expanded samples offering a large FOV of 640 µm x 640 µm, high lateral resolution of ∼566 nm (∼25-30 nm after expansion) and high axial resolution of ∼457 nm, and a fast image acquisition speed of up to 20 fps. We demonstrate the high-speed nanoscale volumetric imaging capabilities of the system by imaging expanded HeLa cells and mouse kidney and brain tissues and comparing it side-by-side with state-of-the-art spinning disk confocal microscopy.

## Results

### Light sheet microscope design and calibration

A key requirement for pan-ASLM imaging of large volumes is the right set of objective lenses that allow for high isotropic resolution, large FOVs, and long WDs, yet can be arranged orthogonally to each other without mechanically interfering. As we are interested in imaging expanded samples which consist mostly of water, we limited our search to objectives that were designed for this medium. We chose a multi-immersion objective (ASI, 54-12-8) with a numerical aperture (NA) of 0.64 (at refractive index n=1.33) and 10 mm WD as our illumination objective (IO) and paired it with a 20x/1.0 NA water-dipping objective (Evident XLUMPLFLN20XW) for detection in an orthogonal geometry (**Figure 1b**). The high NA, low magnification, and long WD (2 mm) of the detection objective allow for high-resolution large-volume imaging. Additionally, since expanded samples can be soft, we oriented the objectives such that the samples are mounted horizontally to minimize distortions by gravitational forces. We use a cylindrical lens to generate the light sheet and a piShaper to ensure uniformity of the intensity across the width of the sheet. As our remote-focusing objective (RFO), we chose a high-NA air objective (Evident, UPLXAPO20X/0.8 NA) which has an angular aperture larger than the IO to satisfy the remote focusing condition [24, 25]. To maximize the FOV, we take advantage of the full chip of a 10-Mpixel camera (Kinetix, Teledyne Photometrix, 3.2k×3.2k pixels, 6.5 µm × 6.5 µm pixel size). We targeted a 200-nm effective pixel size since it provides a good compromise between a large FOV (640 × 640 µm^2^) and a Nyquist-limited resolution close to the diffraction limit (theoretical diffraction-limited resolution: ∼310-430 nm in the 510-700 nm fluorescence wavelength range). This is achieved using an Evident Super Wide Tube Lens (SWTLU-C) with a clear aperture of 36 mm (to not clip the large FOV) and a 1.6x Evident magnification changer (U-CA).

**Figure 1:**
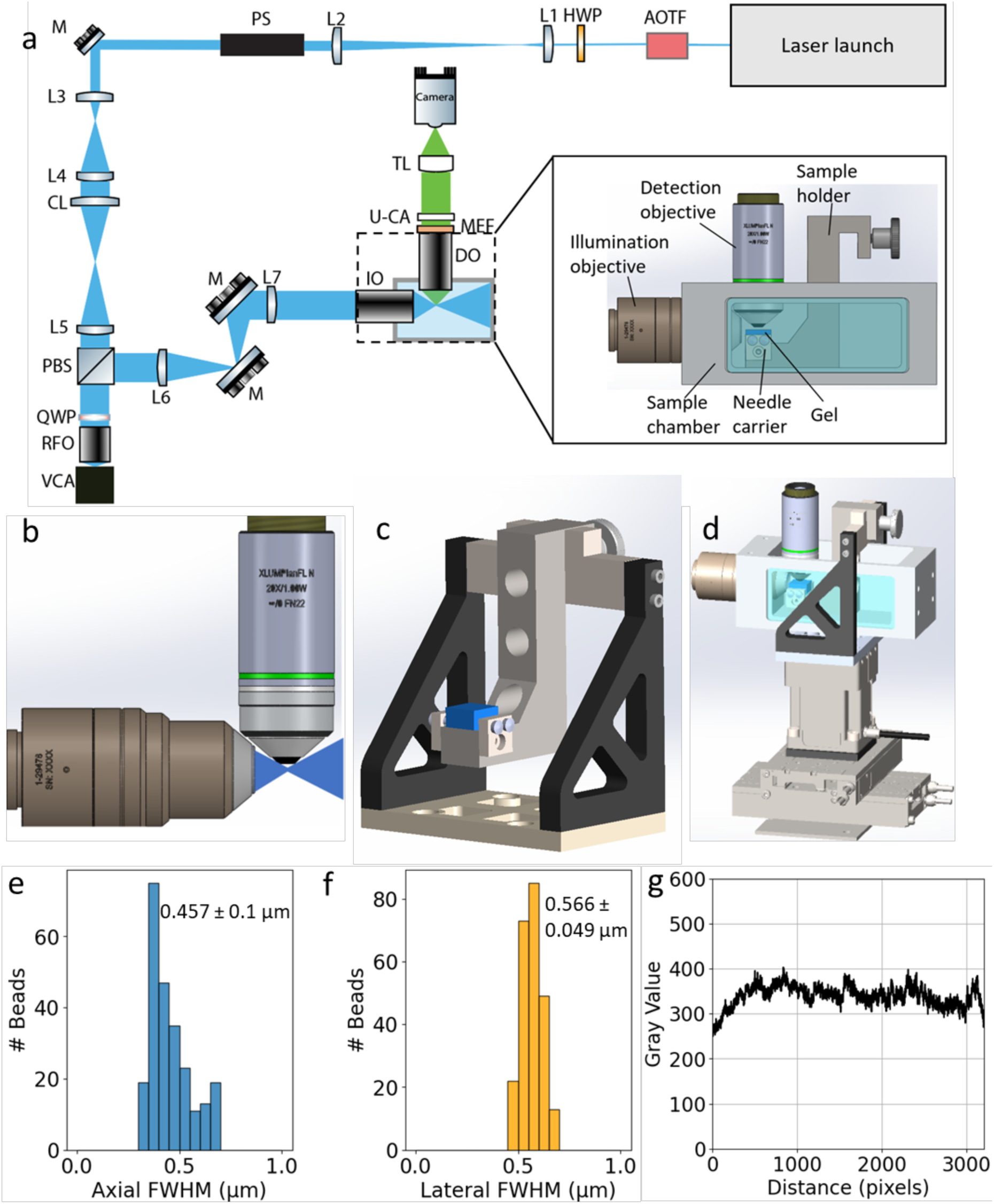
pan-ASLM setup and characterization. **a** Optical layout. AOTF: acousto-optic tunable filter, HWP: half-wave plate, L1-7: lenses, PS: piShaper, M: mirrors, CL: cylindrical lens, PBS: polarizing beam splitter cube, QWP: quarter-wave plate, RFO: remote focusing objective, VCA: voice coil actuator, IO: illumination objective, DO: detection objective, MEF: multiband emission filter, U-CA: U-CA magnification changer lens, TL: tube lens. **b** CAD design illustrating the geometry of the objectives and the aperture angle of the light sheet. **c** Sample holder with the sample shown in blue. **d** Sample holder assembly with XYZ positioning stages. **e, f** Histograms of axial (**e**) and lateral (**f**) FWHM measurements of 100-nm yellow-green beads. **g** Brightness uniformity across FOV for 488-nm excitation.

To filter out the non-focused components of the axially swept light sheet, the sweeping motion induced by the voice coil is synchronized with the rolling shutter of the camera, which at a width of 8 pixels, or ∼1.6 µm, roughly matches the depth of focus of our light sheet [20]. To realize uniform light-sheet properties across the complete FOV, this synchronization needs to be precise to about half the rolling shutter width (i.e. 4 pixels, or 800 nm) which corresponds to ∼0.13% of the FOV. This high relative precision requires accounting for inherent nonlinearities in the voice coil scan and optics by careful calibration (**Supplementary Figure 1**). We first generate a calibration curve that correlates the voltage controlling the voice coil with the position of the light sheet focus as observed by the camera through the IO. For this purpose, the cylindrical lens in the beam path is replaced by a spherical lens of the same focal length, which creates an easily visualizable focus. The voice coil is then parked at ∼100 discrete positions about equidistantly spread across the FOV and the corresponding focus positions are determined from the camera images by finding the brightest pixel. We thus obtained a calibration curve of focus position versus voice coil voltage, which can then be inverted to generate a series of voltage values to be applied to the voice coil to synchronize the light sheet focus with the rolling shutter. This calibration method is suitable for low scan rates of 1 Hz where inertia of the voice coil motion is negligible. For higher imaging speeds, however, inertia prevents the voice coil to precisely follow an applied voltage scan pattern resembling high-frequency triangular or sawtooth functions required for the axial scan of the light sheet and the corresponding backscan. This causes a deviation of the actual light sheet focus position from the corresponding rolling shutter position which needs to be corrected for. To enable higher scan rates, we used a global optimizer to determine sets of parameters, each for a particular frame rate, of a third-order polynomial function used for generating the voice coil voltage signal that best synchronizes the light sheet focus with the rolling shutter. Image analysis is the basis for error evaluation, which is minimized through the optimization scheme. This process allowed for the calibration of the voice coil motion for high scanning speeds of up to 20 Hz. More details of the voice calibration process can be found in **Supplementary Note 1**.

To characterize the resolution of the system, we imaged green fluorescent beads embedded in 1% agarose. We measured the Full-Width-at-Half-Maximum (FWHM) of each bead (n=243) and found 0.457 ± 0.1 µm axial resolution and 0.566 ± 0.049 µm lateral resolution (**Figure 1e,f**). This nearly isotropic resolution makes the instrument ideal for imaging tissue samples in 3D.

The brightness uniformity across the FOV was measured by taking a line profile averaged in the scanning-direction of the swept light sheet and showed that the brightness dropped only by ∼25% towards the edge of the FOV (**Figure 1g**).

To test the compatibility with expanded samples, we imaged expanded and pan-stained HeLa cells (**Figure 2**), revealing subcellular features at the nanoscale such as mitochondria cristae (**Figure 2b**), the substructure of nucleoli (**Figure 2c**), centrioles (**Figure 2d, g**), nuclear pore complexes (**Figure 2h**), and putative annular lamellae (**Figure 2i**) at excellent contrast.

**Figure 2:**
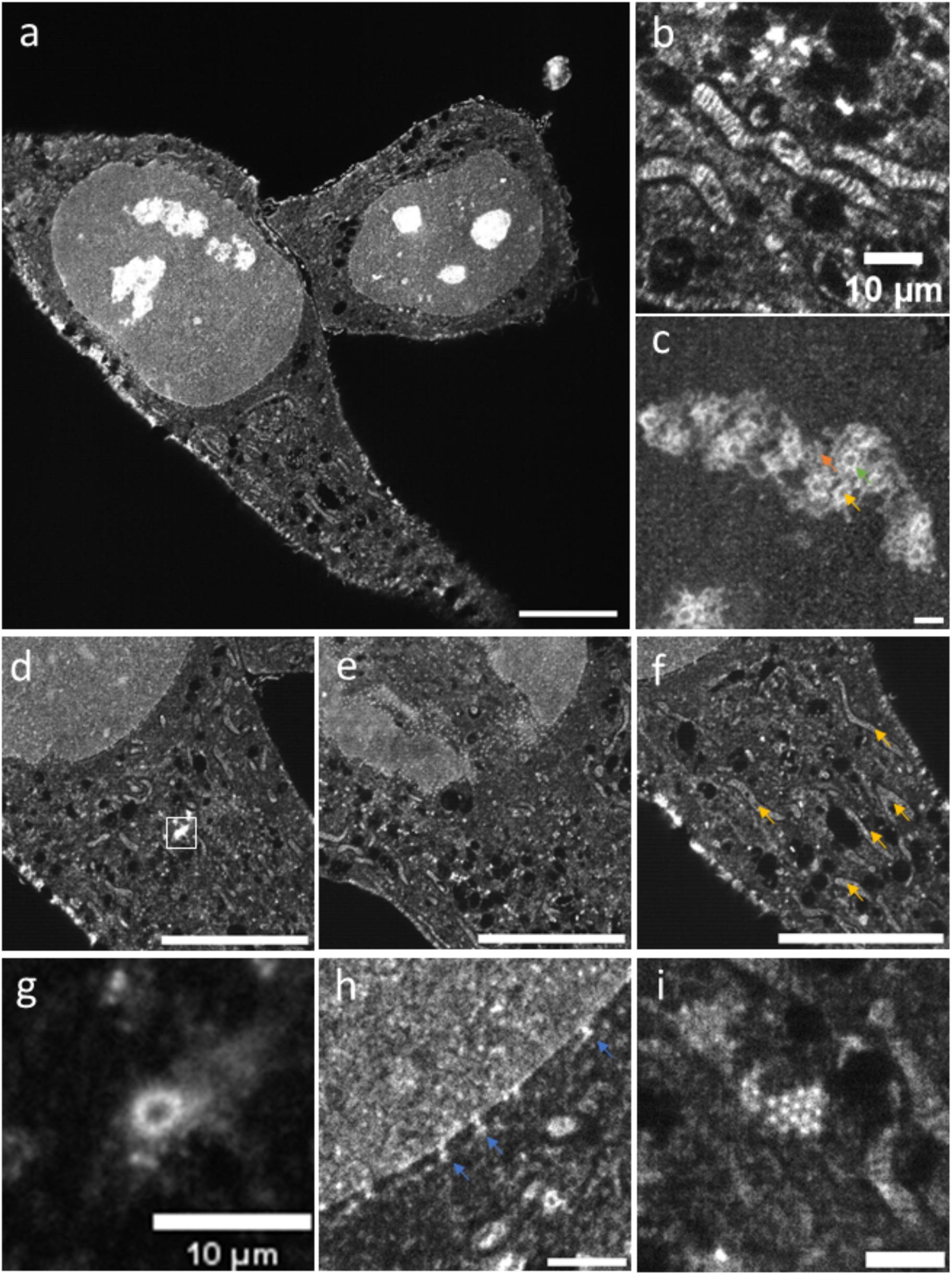
pan-ASLM images of pan-expanded HeLa cells. **a** A single FOV image of two pan-expanded HeLa cells stained with Atto 488 NHS ester. **b** Zoomed-in view of mitochondria showing clearly resolvable cristae. **c** Zoomed-in view of nucleolus showing dense fibrillar component (green arrow), fibrillar center (yellow arrow), and granular component (orange arrow). **d** Zoomed-in view of HeLa cell from the same dataset as (**a**). **e** Zoomed-in view showing nuclear pore complexes. **f** Zoomed-in view showing mitochondria (orange arrows). **g** Zoomed-in and contrast-adjusted view of the white box in (**d**) showing a centriole. **h** Zoomed-in view of the nuclear envelope showing nuclear pore complexes (blue arrows). **i** Zoomed-in view of putative annular lamellae in the cytoplasm. Scale bars are not corrected for the expansion factor. Scale bars: (**a, d-f**) 100 μm, (**c, h, i**) 10 μm.

### Comparison with spinning disk confocal microscopy

To evaluate the imaging performance of our pan-ASLM instrument, we benchmarked it against an Andor Dragonfly 600 mounted to a Nikon Ti-2 inverted microscope stand, a state-of-the-art spinning disk microscope which we frequently rely on in pan-ExM applications due to its excellent combination of 3D resolution and imaging speed [13]. Because the spinning disk microscope image quality is optimized for high-magnification objectives (see above), we used a 60x/1.2 NA water immersion objective for the comparison. We imaged the same 16x expanded mitotic HeLa cell pan-stained with CF 568 NHS ester on both instruments which allowed us to directly compare their imaging performance **(Figure 3)**. Both instruments clearly resolve the cristae inside mitochondria, a hallmark feature of super-resolution microscopy. However, zooming into XZ and YZ views of a mitochondrion reveals that the pan-ASLM, due to its ∼1.7x superior axial resolution (457 nm vs. 800 nm), resolves the 3D shape of the organelle much better than the spinning disk system. Interestingly, the slightly worse XY resolution (caused by the lower NA of 1.0 in pan-ASLM vs. 1.2 for the spinning disk) and larger pixel size (200 nm vs. 108 nm) do not substantially limit our capability to clearly distinguish the cristae in the mitochondrion, supporting the notion that a good isotropic resolution is often more important when imaging complex 3D structures than maximizing the resolution in some directions at the cost of others. In addition, **Figure 3a** illustrates the ∼7-fold large FOV of pan-ASLM (640 x 640 µm^2^) compared to the spinning disk system (∼251 x 251 µm^2^). Furthermore, the free working distance of the pan-ASLM of 2 mm is ∼7-fold larger than that of the 60x/1.2 NA objective (280 µm). In addition, pan-ASLM achieves 20 fps imaging while the spinning disk instrument achieves ∼3.3fps for high quality datasets, corresponding to ∼15x faster imaging speed of the pan-ASLM (183 Mvoxels/sec vs ∼12 Mvoxels/sec).

**Figure 3:**
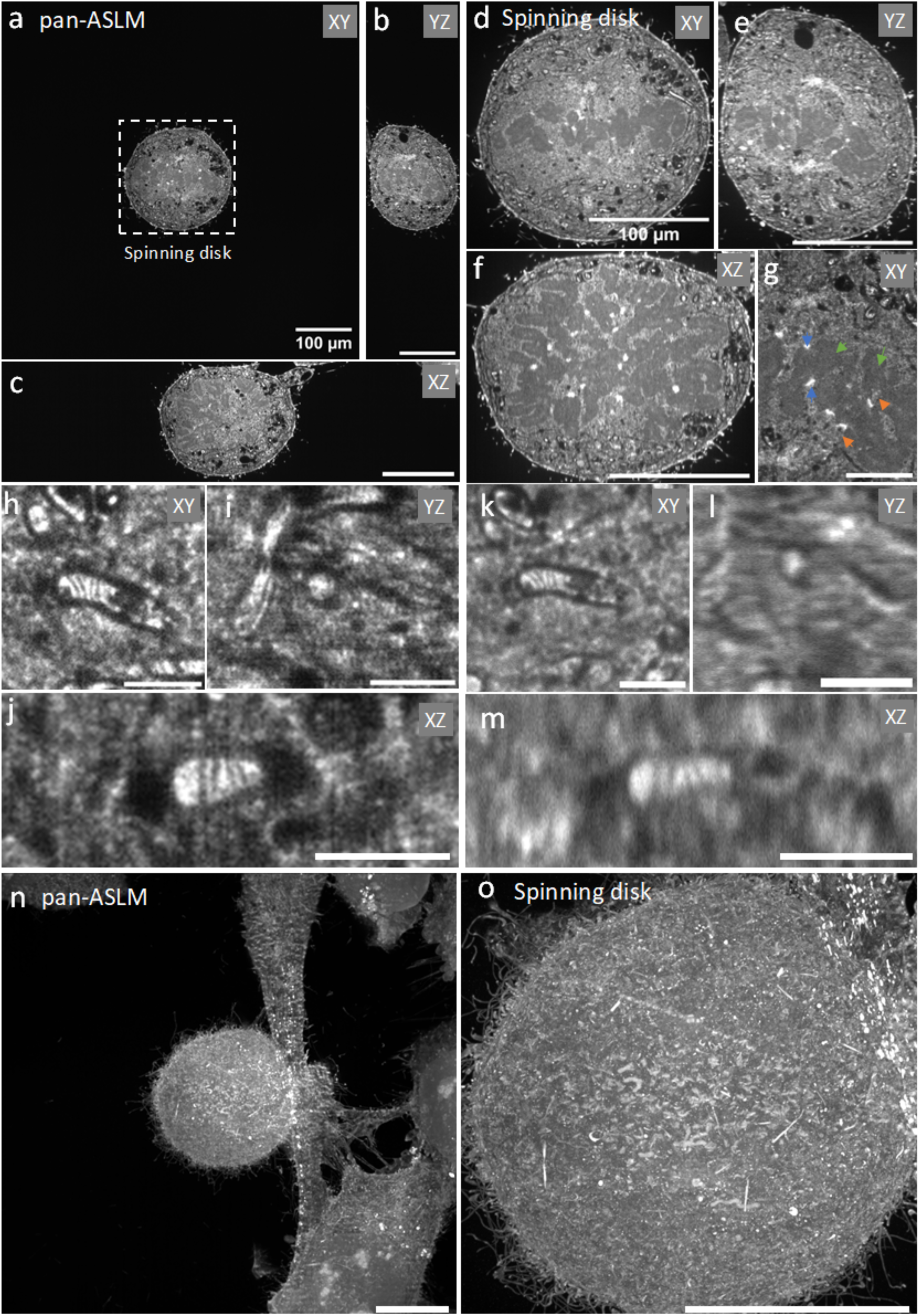
Comparison between pan-ASLM and spinning disk microscopy images. **a-c** XY (**a**), YZ (**b**), and XZ (**c**) optical slices of the 3D pan-ASLM data set of a pan-expanded mitotic HeLa cell stained with CF 568 NHS ester. The box in (a) shows the FOV of the spinning disk microscope with a 60x/1.2 objective. **d-f** XY (**d**), YZ (**e**), and XZ (**f**) slices of the 3D data set of the same mitotic cell imaged on the spinning disk microscope with a 60x/1.2 objective. **g** Zoomed-in region of the mitotic cell imaged with pan-ASLM showing kinetochores (blue arrows), microtubules (orange arrows) and chromosomes (green arrows). **h-j** Zoomed-in XY (**h**), YZ (**i**), and XZ (**j**) slices of zoomed-in volume showing a mitochondrion imaged with the pan-ASLM. **k-m** XY (**k**), YZ (**l**), and XZ (**m**) slices of the corresponding volume imaged with spinning disk microscopy. **n** Maximum Intensity Projection (MIP) of a mitotic cell and neighboring cells imaged with pan-ASLM. **o** MIP of the same mitotic cell imaged with spinning disk microscopy (FOV has not been cropped). Scale bars are not corrected for the expansion factor. Scale bars: (**a-g, n, o**) 100 μm, (**h-m**) 10 μm.

### High-speed and multicolor imaging of pan-expanded HeLa cells

To test the high-speed and multicolor imaging capabilities of the microscope, we imaged pan-expanded HeLa cells at different speeds ranging from 2 fps to 20 fps (**Figure 4**). Nanoscale details of organelles such as mitochondria cristae, nuclear pore complexes and kinetochores, all revealed by the NHS ester pan-stain, can be discerned equally well from images recorded at 2 fps and 20 fps. 3-color data collection is demonstrated by imaging, in addition to the pan-stain (CF 568 NHS ester) in the orange channel, the DNA (SYTOX Green) in the green spectral range and the outer mitochondrial membrane (anti-TOM20 labeled with Atto 647N) in the far-red range of the spectrum. The large FOV allows for imaging multiple cells in a single FOV (**Figure 4f**) that would have otherwise required a mosaic of multiple tiles if imaged on the spinning disk microscope. We were able to image the whole 3D volume of expanded cells in less than a minute as shown in **Figure 4l, m**.

**Figure 4:**
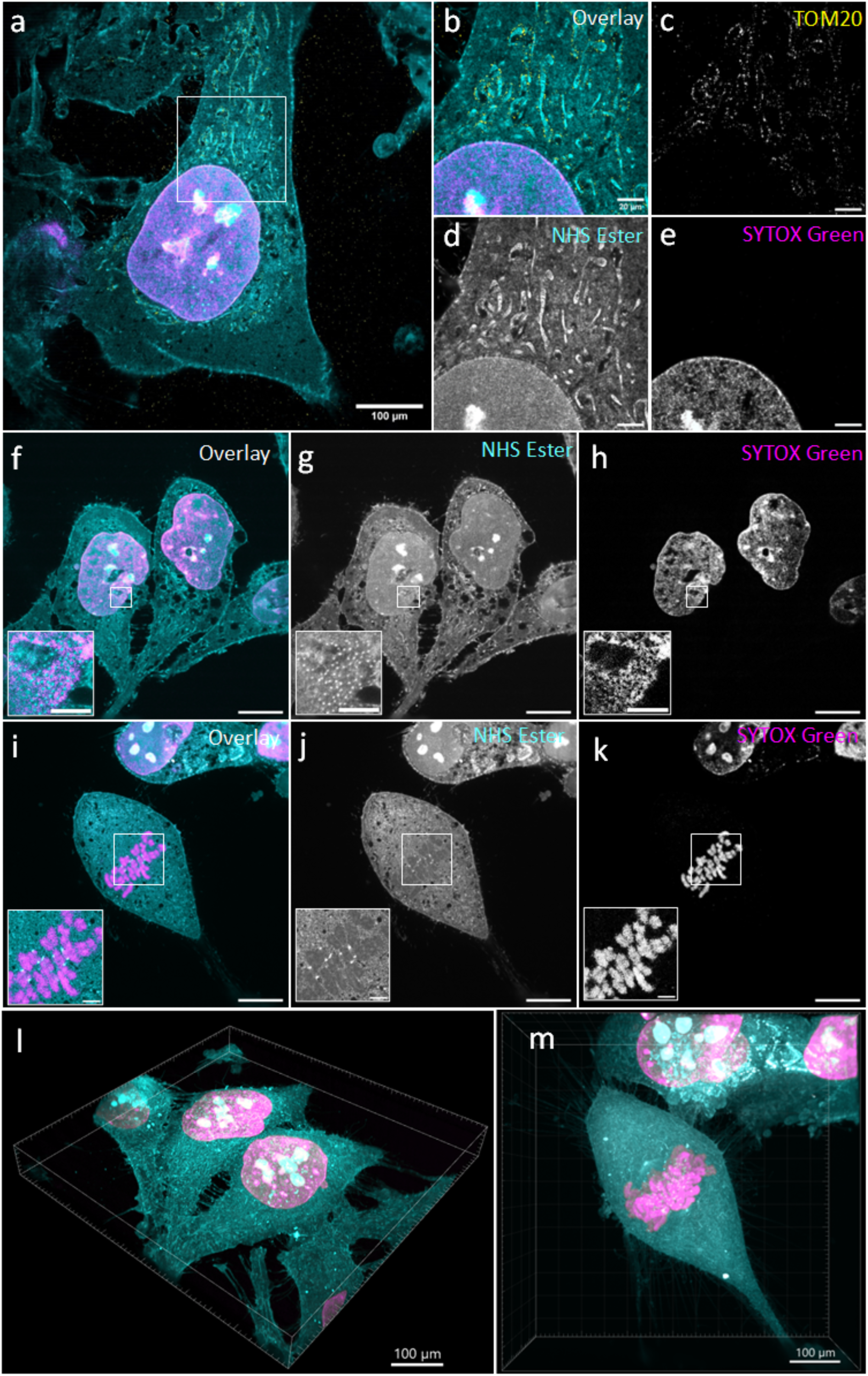
High-speed and multicolor imaging of pan-expanded HeLa cells. **a** pan-ASLM overlay image of a 3-color pan-expanded HeLa cell labeled with CF 568 NHS ester (shown in cyan), SYTOX Green (shown in magenta) and anti-TOM20 (Atto 647N; shown in yellow) imaged at 2fps. **b-e** Zoomed-in overlay (**b**) and single Anti-TOM20 channel (**c**), CF 568 pan-stain channel (**d**), and SYTOX Green (**e**) channel images of the area shown in the white box in (**a**). **f-h** 2-color pan-ASLM overlay (**f**) and individual channel (**g**, **h**) images of pan-expanded interphase HeLa cells recorded at 10 fps (76 ms acquisition and 24 ms flyback time). The insets show zoomed-in views of nuclear pore complexes. **i-k** 2-color pan-ASLM overlay (**i**) and individual channel (**j**, **k**) images of a pan-expanded mitotic HeLa cell recorded at 20 fps (26 ms acquisition and 24 ms flyback time). The insets show zoomed-in views of chromosomes and kinetochores. **l** 3D view of the data set shown in (**f**). **m** MIP of the data set shown in (**i**). Scale bars are not corrected for the expansion factor. Scale bars: (**a, f-k**) 100 μm, (**b-e**) 20 μm, (insets **f-k**) 20 μm.

### Large-field-of-view imaging of pan-expanded mouse kidney and brain tissue

To validate the system for large-FOV imaging in tissue samples, we imaged pan-expanded mouse kidney tissue and applied 2D tiling. We used the open-source software BigStitcher [35] for stitching the images, resulting in 3.86 mm x 2.16 mm large data sets of expanded tissue sections as shown in **Figure 5a**. Zooming into these data sets revealed easily resolvable details at the nanometer scale such as the mitochondria cristae (**Figure 5e, f, j**), podocyte foot processes (**Figure 5o, p**) in the glomerulus, and the brush border (**Figure 5h, i**) in the proximal tubule. The isotropic resolution of the system is also illustrated in **Figure 5d, f, n, p** where the XY views and XZ views show an image quality indistinguishable by eye.

**Figure 5:**
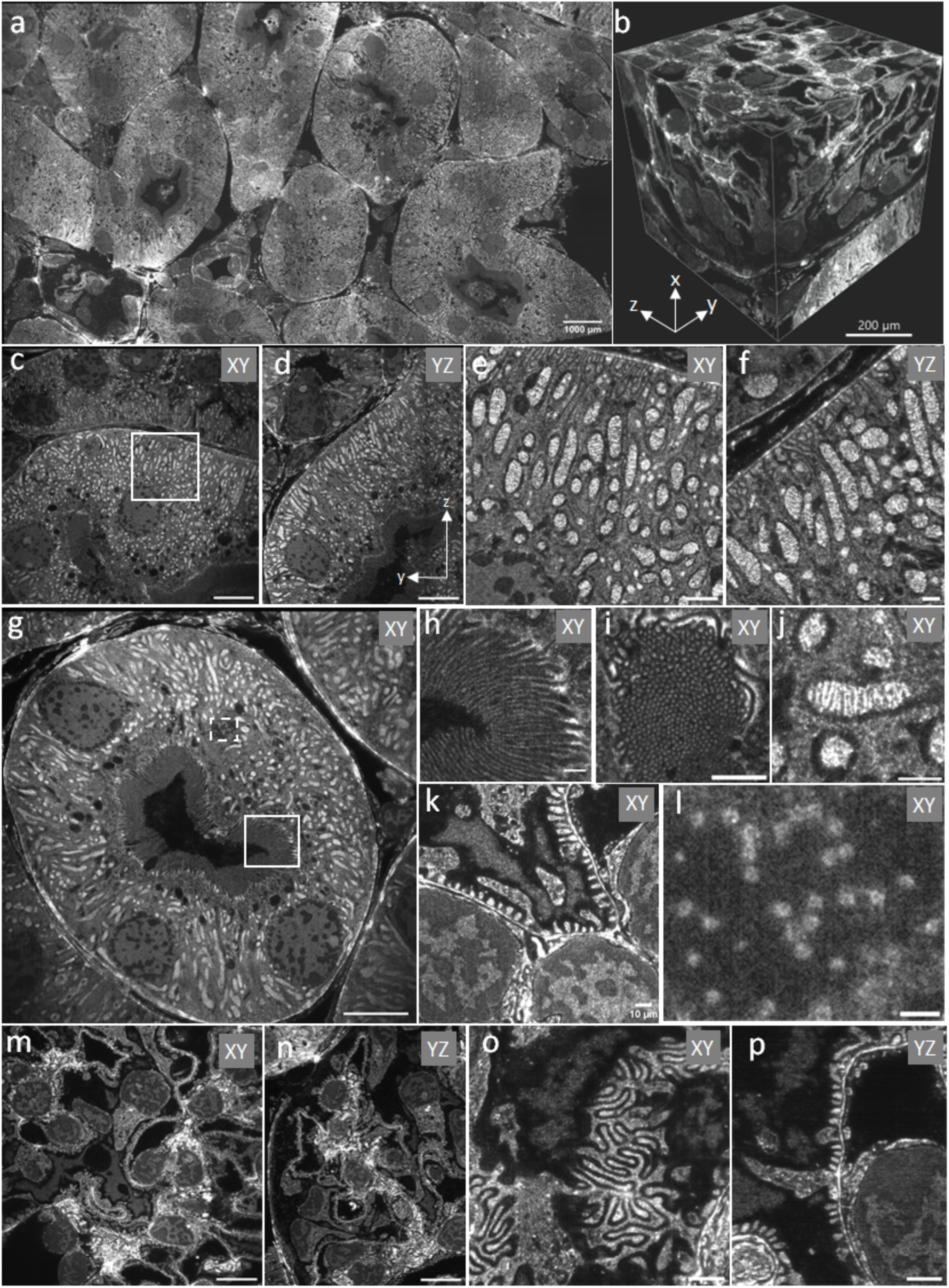
Large-FOV imaging of pan-expanded mouse kidney tissue. **a** Stitched pan-ASLM image of a 3.86 mm x 2.16 mm large pan-expanded mouse kidney tissue area pan-stained with Atto 488 NHS ester. **b** 3D volume of a glomerulus. **c** A single XY slice of a proximal tubule. **d** YZ view of the same region as shown in (**c**). **e** Zoomed-in view of the white box shown in (**c**). **f** YZ view of the same region as shown in (**e**). **g** A single XY slice of another proximal tubule. **h** Zoomed-in view of the solid white box drawn in (**g**) showing the brush border. **i** XY slice through Z-oriented microvilli of the brush border. **j** Zoomed-in view of a mitochondrion. **k** Zoomed-in view of podocyte foot processes found in the glomerulus. **l** Zoomed-in view of the dashed white box in (**g**) showing nuclear pore complexes and their ring-like structure. **m, n** XY (**m**) and YZ (**n**) slices through the glomerulus shown in (**b**). **o** Zoomed-in view of the glomerulus showing podocyte foot processes. **p** YZ view of the region shown in (**o**). Scale bars are not corrected for the expansion factor. Scale bars: (**c, d, g, m, n**) 100 μm, (**e, i**) 25 μm, (**f**, **h, j, l**) 10 μm, (**l**) 5 μm, (**o, p**) 20 μm.

To test the large-volume imaging capabilities of the microscope, we further imaged a 6.11 mm x 3.07 mm large area of a pan-expanded mouse cortex sample (**Figure 6a**) over a depth of 1.2 mm (**Figure 6b**). The image quality across the tissue section is preserved as shown in **Figure 6c-e**. The long WD of the IO reduces the need for sectioning expanded samples and substantially simplifies imaging large volumes. The pan-ExM technique allows for visualization of individual synapses [13], and the isotropic resolution of the system allows for accurate representation of synapses independent of their orientation in 3D (**Figure 6k-n**).

**Figure 6:**
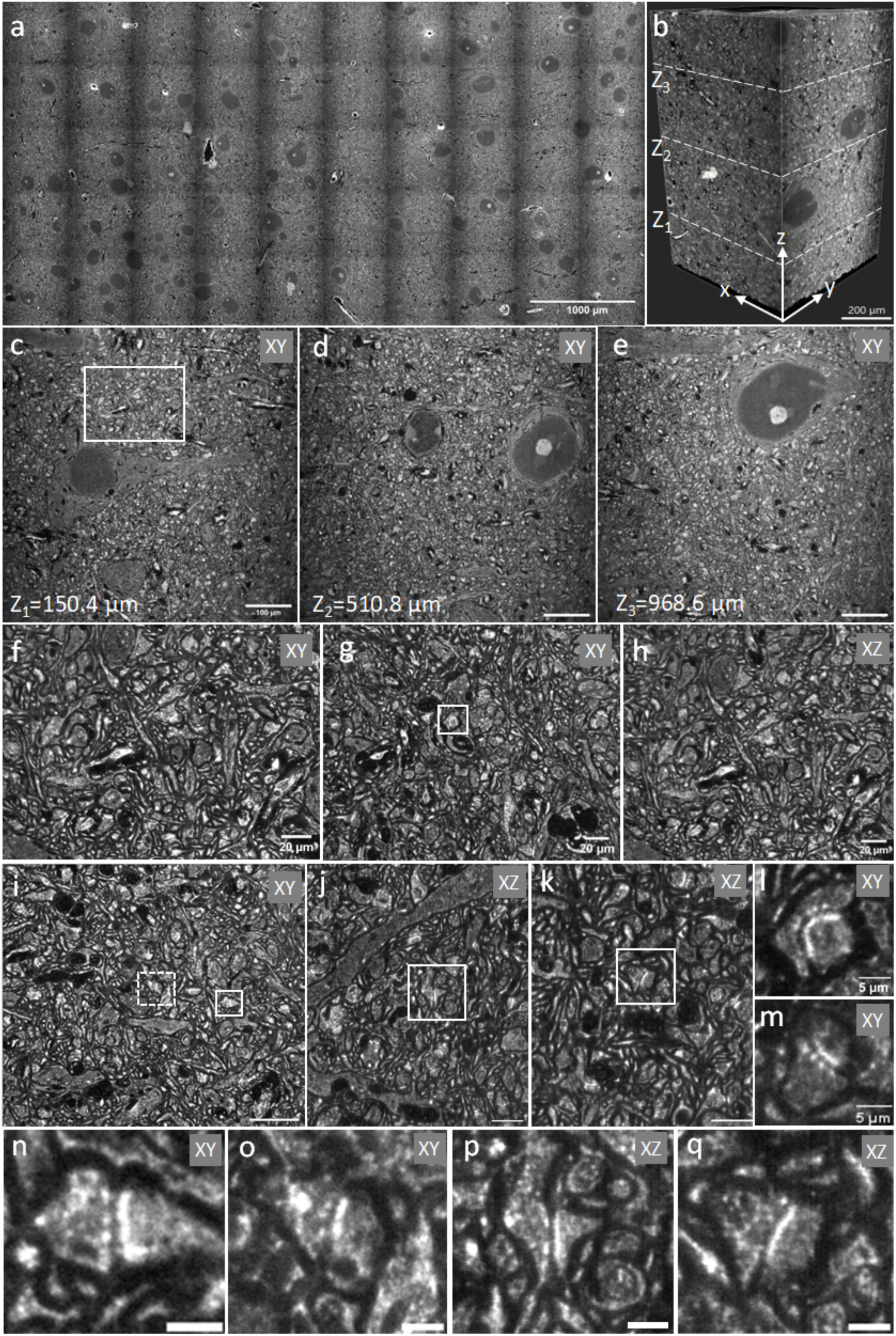
Large-volume imaging of pan-expanded mouse cortex. **a** Stitched pan-ASLM image of a 6.11 mm x 3.07 mm large area of a pan-expanded mouse cortex with NHS ester pan-staining. **b** 3D volume of a pan-expanded ∼1.2-mm thick mouse cortex. **c-e** Single XY slices of the volume shown in (**b**) at Z=150.4 µm (**c**), Z=510.8 µm (**d**), and Z=968.6 µm (**e**). **f** Zoomed-in view of the white box in (**c**). **g** Zoomed-in XY slice of a region from (**b**). **h** XZ slice of the same region as shown in (**g**). **i** Another zoomed-in XY slice of a region from (**b**). **j, k** Two XZ slices of the same region as shown in (**i**). **l** Zoomed-in view of the synapse in the white box shown in (**g**). **m** Zoomed-in view of a synapse from (**b**). **n, o** Zoomed-in views of the solid (**n**) and dashed (**o**) white boxes in (**i**) showing individual synapses. **p** Zoomed-in view of the white box in (**j**). **q** Zoomed-in view of the white box in (**k**). Scale bars are not corrected for the expansion factor. Scale bars: (**c-e**) 100 μm, (**f-k**) 20 μm, (**l-q**) 5 μm.

## Discussion

We have built with pan-ASLM a new LSFM that enables high-throughput imaging of expanded samples, cutting down imaging times from hours to minutes when compared to a conventional confocal microscope. pan-ASLM provides high isotropic resolution and image acquisition speeds of up to 20 fps. To our knowledge, the microscope provides the fastest speed and highest FOV-to-resolution ratio among current high-resolution ASLM variants [25, 34]. With large-volume ExM applications rapidly emerging, pan-ASLM meets the growing need for high-throughput, large-FOV, long-WD microscopes that perform at the resolution level of the best confocal microscopes.

At the obtained ∼500 nm isotropic resolution, ∼20-fold expanded samples are resolved at effectively 25 nm - fully sufficient to resolve mitochondria cristae, microvilli of the brush border in proximal tubules of the kidney, and the synaptic cleft between neurons in the mouse brain. In cases where additional resolution should be required, the images could be deconvolved to gain a factor of ∼1.5 of additional resolution improvement, as has been shown in other ASLM publications [25, 33].

Further modifications to the current microscope can be made to improve its performance. While we currently use a camera with 10 Mpixels, the available pixel number is steadily increasing with cameras featuring well beyond 20 Mpixels already on the market. These cameras offer the potential to increase the FOV even further and/or reduce the effective pixel size to get closer to the diffraction-limited performance of the optical detection system. In addition, future objectives with longer WDs while maintaining high NAs will increase the sample volume accessible for imaging without compromising the resolution. Our imaging speed is currently limited by the speed at which we can scan the voice coil in synchrony with the rolling shutter of the camera. Optimizing the mechanics of this motion using more responsive actuators should be able to raise the frame rate to the maximum camera frame rate. With each camera generation offering higher frame rates, high-power lasers becoming more readily available, and more photostable dyes and imaging buffers being invented, the potential for future improvements of ASLM imaging speeds is large.

With the increase in data acquisition throughput, the spotlight moves to automated image processing and quantification. We have recently developed an automated image segmentation and quantification approach for pan-ExM data sets acquired with spinning disk confocal microscopes [14]. Given that the image quality we have demonstrated here surpasses that of spinning disk confocal microscopes, we anticipate that this deep-learning based approach can be readily adopted to our large pan-ASLM data sets. This combination is particularly attractive in applications such as the connectomics field, where large-scale mapping of neuronal circuits and synaptic proteins across neurons over large volumes is essential. pan-ExM and related ExM approaches have recently been demonstrated to provide sufficient contrast for neuronal tracing [13, 36] and have the potential to replace electron microscopy as the imaging method of choice for large-scale connectomes. Connectomics represents, however, just one of many biomedical questions that will benefit from large-volume, high-resolution imaging as provided by pan-ASLM and we anticipate a wealth of applications exploring biological heterogeneity across large numbers of cells and tissue volumes.

## Methods

### Light sheet microscope

A parts list is provided in **Supplementary Table 1**. Beams of four lasers with wavelengths of 642 nm (MPB Communications, 2 W), 595 nm (MPB Communications, 500 mW), 561 nm (MPB Communications, 500 mW), and 488 nm (Coherent OBIS 488, 150 mW) are combined with dichroic mirrors into a common beam path and pass through an acousto-optical tunable filter (AOTF; AOTFnC-VIS, AA Optoelectronic) for power adjustment. A piShaper (#12-644, Edmund Optics) is used to generate a uniform intensity profile after the beams are expanded to the required diameter by a 4-f telescope (f=50 mm, AC254-50-A; f=300 mm, AC254-300-A). The beam is further expanded by a factor of 3 before entering a cylindrical lens (f=100 mm; ACY254-100-A; Thorlabs) which focuses the beam into a light sheet which is imaged by a 150-mm focal lens achromat (AC254-150-A, Thorlabs) and the remote focus objective (RFO) (Evident, UPLXAPO20X/0.8 NA) into the RFO’s focal plane. A small mirror (BB03-E02, Thorlabs), mounted to a voice coil (LFA 2010, Equipment Solutions), reflects the beams back through the RFO. After passing through a lambda/4 plate on the way into and out of the RFO, the laser light is reflected by a polarizing beam splitter cube (CCM1-PBS251/M, Thorlabs) towards the illumination objective (IO) (ASI, 54-12-8). A 4-f system consisting of two achromats (f=150 mm, AC254-150-A; f=200 mm, AC254-200-A) combined with a periscopic mirror arrangement images the back pupil plane of the RFO into the back pupil of the IO, resulting in the light sheet, axially swept by the voice coil motion, being projected into the focal region of the IO. A 20x/1.0 NA water-dipping objective (XLUMPLFLN20XW, Evident) is used for detection. A multiband emission filter (ZET405/488/561/640mv2, Chroma) above the objective is used to filter out scattered laser light. A large-FOV tube lens (SWTLU-C, Evident) and a 1.6x magnification changer (U-CA, Evident) are used to form the image on the sCMOS camera chip (Kinetix, Teledyne Photometrics, 3.2k×3.2k pixels, 6.5 µm × 6.5 µm pixel size). A custom-designed sample chamber and sample holder are used to hold expanded gels in a horizontal orientation. A flat-top Z-stage (Physik Instrumente, L-306.011112), two XY linear stages (Physik Instrumente, V-508.231) and a multi-axis stage controller (G-901.R3197) are used for XYZ positioning of the sample. In addition, the microscope uses a piezo objective actuator (Thorlabs, PIA13) for Z-focusing. The custom-written LabVIEW software allows for automated recording of large, tiled data sets to cover volumes as large as ∼10×10×2 mm^3^. In order to synchronize the rolling shutter of the camera with the voice coil, we use a NI DAQ card (PCIe-6323) that sends signals to both the voice coil and the camera.

### Acquisition software

We use custom-written LabVIEW software to control the microscope. The acquisition code and voice coil calibration code are available on GitHub at https://github.com/lasselpk/pan-ASLM/.

### Sample mounting

We use a custom-designed sample mount to mount the gels for imaging. 200 µL of 0.1% Poly-L-lysine solution (P8920, Sigma) are added on the sample holder. After letting it dry on a heat block for 10 minutes, the gel is placed onto the mount. This will attach the gel to the mount and prevent drift during imaging. We use ultra-thin walled needles (34G, AD Surgical) to mount the gels by gently pushing the needles on either side into the needle carriers.

### Resolution measurement

To measure the system’s point-spread function (PSF), we embedded 100-nm diameter yellow-green beads in a 1% agarose solution. After pouring the solution onto a petri dish and letting it cool, we used a razor blade to cut a rectangular (∼10 mm x 10 mm) section and placed it on the sample holder for imaging. We used a 488-nm laser to excite the beads. The sample was imaged with a Z-step size of 200 nm and 4 pixels rolling shutter scan width. We then calculated the PSF by measuring the FWHM from the beads’ line profile using PSFj, a software tool designed to measure PSFs of different microscope systems [37]. We then plotted the data using modified python code from Valdimirov et al. [38].

### Sample Preparation

HeLa cell samples were prepared as described by M’Saad, et al. [12]. Mouse brain and kidney tissue samples were prepared as described by M’Saad, et al. [13] and Tian, et al. [14].

## Code availability

The microscope control software is available at https://github.com/lasselpk/pan-ASLM/.

## Acknowledgements

We thank Drs. Reto Fiolka and Kevin Dean for advice during the early phase of microscope development. We also thank Dr. Lin Shao for help with LabVIEW programming and comments on the manuscript. L.P.A. is supported by a NERD grant from the Novo Nordisk Foundation (NNF20OC0059893). J.G. acknowledges support by the NIH (R43DK137685; R44MH128999). J.B. acknowledges support by the Wellcome Leap foundation.

## Author Contributions

H.T.M. and J.B. designed the microscope. H.T.M. built the microscope. M.L. designed the sample holder assembly. H.T.M., L.P.A., and J.R. wrote the custom-written LabVIEW code for the microscope control. S.Z. prepared pan-expanded HeLa cell samples. J.G. prepared pan-expanded mouse brain and kidney tissue. Y.T. prepared pan-expanded mouse brain tissue. J.B. supervised the project. H.T.M. and J.B. wrote the manuscript. All authors commented on the manuscript.

## Competing Interests

J.B. is a co-founder of panluminate, Inc. J.G. is an employee of panluminate, Inc.

