## Supplemental Information for "Pan-ASLM: a high-resolution and large field-of-view light sheet microscope for Expansion Microscopy"

### Supplementary Note 1

#### Voice coil calibration for high-speed imaging

In pan-ASLM, we use a voice coil actuator for axial sweeping of the light sheet. Replacing the cylindrical lens by a spherical lens forms a beam focus that can be used to visualize the synchronization between the voice coil actuator and the rolling shutter of the camera. Sending a sawtooth waveform to the voice coil, without using the rolling shutter mode, results in an axially uniform pattern that forms across the field of view (**Supplementary Figure 1a**), showing that the remote focusing allows for translation-invariant focus quality. However, in the rolling shutter mode, the line profile degrades as the axial sweeping and the rolling shutter get out of synch for most of the voice coil travel range. This shows that a careful calibration of the voice coil is necessary to achieve synchronization between axially sweeping the light sheet and the rolling shutter of the camera.

To calibrate the voice coil, we first generate a list of voltage values within the range of the FOV and acquire images for the beam focus corresponding to each voltage value. We use Picasso [1] to fit a gaussian function to the brightest pixel position in each image. For a set image acquisition time, we plot the position of the voice coil as a function of time, and we obtain a calibration curve after converting it into a voltage-over-time function with which the voice coil can be controlled (**Supplementary Figure 1b**). We next characterize the calibration curve by fitting a third-order polynomial function. After this calibration, we observe a line profile mostly uniform in the axial sweeping direction across the field of view. However, at the bottom of the field of view, corresponding to the start of the sweeping motion, a ringing pattern can be observed. This ringing can be eliminated by adding flyback time to the function controlling the voice coil motion. This calibration method is suitable for low scan rates of 1 Hz.

To enable higher scan rates, we implemented an automated calibration routine for the voice coil scanner using a global optimizer. The global optimization algorithm minimizes an error function, *f(x)* by finding *x*, which is subject to boundary constraints min ≤ *x* ≤ max set by the user. Briefly, based on the user inputs of minimum and maximum boundaries for each polynomial coefficient used to model the calibration curve, the algorithm sends a signal to the voice coil and calculates an error based on the line profile. The error is calculated by first binning the image (2x16) and then cropping the image to 200x200 pixels. We then fit a gaussian function to each row of pixels and calculate the FWHM (**Supplementary Figure 2a**). The sharper the line, the smaller the FWHM. The mean of the FWHM distribution for each image (line profile) can be calculated and used as an error value corresponding to each line profile. The polynomial coefficients are updated until a lower error (mean) value is achieved, which represents a uniform line profile across the field of view. In the present setup, the scanning speed in pan-ASLM is limited by the available laser power (as lower exposure times require higher signal) and by the flyback time as we need to allow some time for the voice coil to equilibrate before subsequent scanning.

#### Supplementary Note 2

#### pan-ASLM imaging volume and sample thickness

Based on the working distance of the detection objective and the sample chamber size, pan-ASLM is able to image a volume of 10x10x2 $mm^3$. However, this volume can be reduced if the sample is less than ~5 mm thick. The reason for this limitation is that the sample holder clips the light sheet when imaging too close to the sample holder, which happens when one images deep into a relatively thin (<2-mm thick) gel both in the X and Z directions (**Supplementary Figure 7**).

#### Supplementary Figures

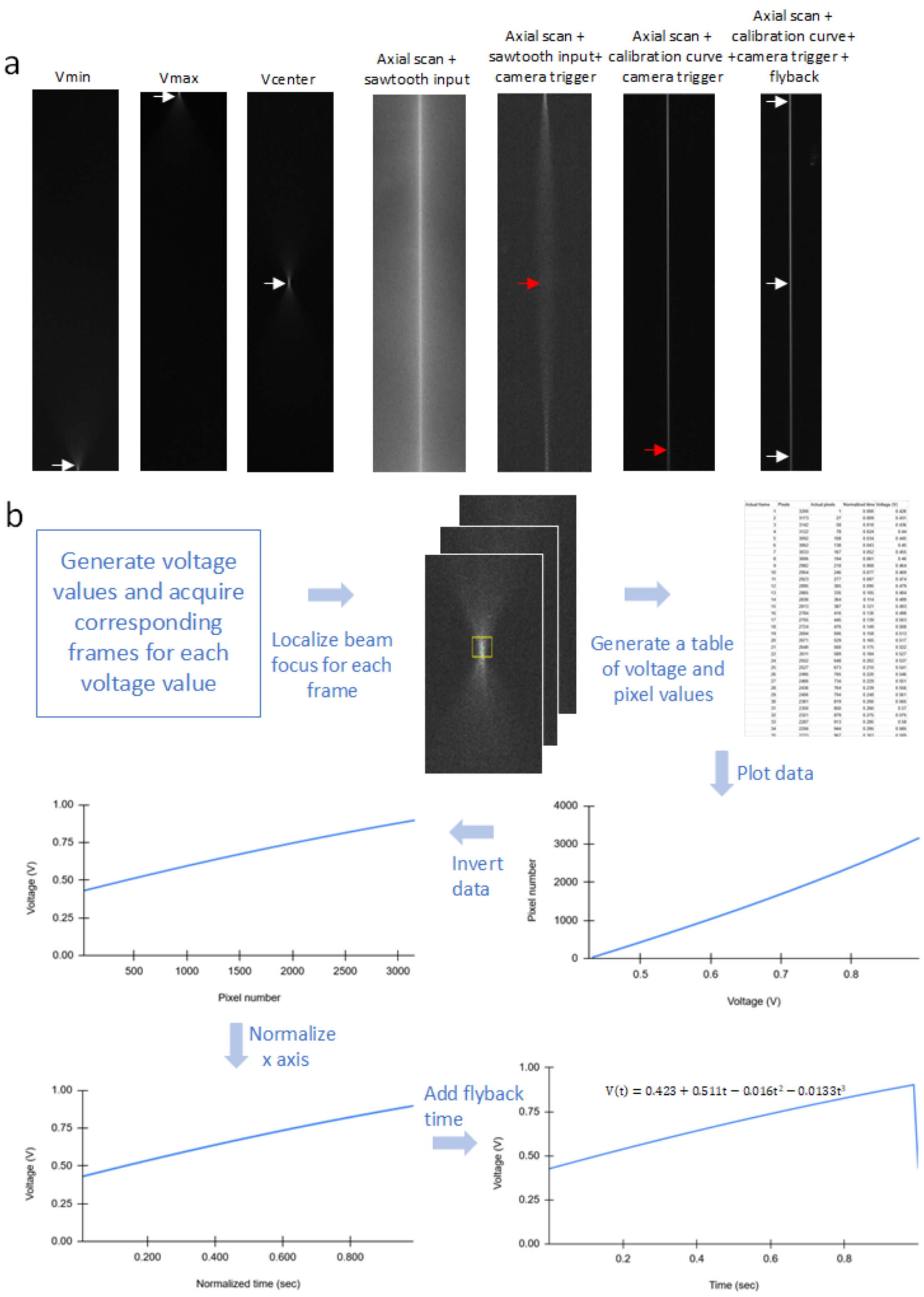


**Supplementary Figure 1: Calibration of voice coil actuator for 1 fps frame rate. a** (from left to right) Beam focus position corresponding to different voltage values, $V_{min}$, $V_{max}$ and $V_{center}$, axial scan without the rolling shutter effect, axial scan with sawtooth voltage input and camera trigger illustrating the lack of synchronization between the voice coil motion and the rolling shutter, axial scan with calibration showing nearly uniform line across the FOV except for a ringing pattern at the bottom of the image, same with flyback time added to the calibration curve resulting in a uniform line across the complete scan range. **b** Process of calibrating the voice coil actuator for 1-Hz scan rate.

a

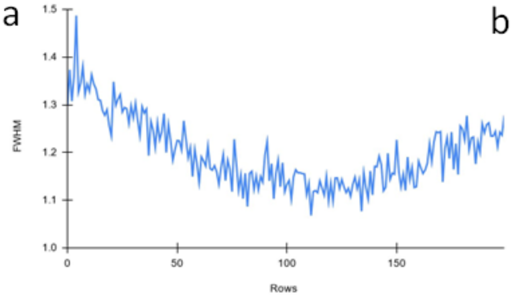


b

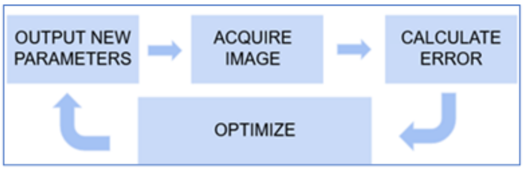


c

| Frame rate | Polynomial function representing calibration curve | FOV (pixels) | Voxel rate (Megavoxels/sec) |
|---|---|---|---|
| 1 fps | $0.423+0.511t-0.016t^2-0.0133t^3$ | 3200 x3200 | 10.2 |
| 5 fps | $0.429+0.53t-0.016t^2-0.0133t^3$ | 3200x3155 | 50.5 |
| 10 fps | $0.434+0.567t-0.0163t^2-0.0137t^3$ | 3200x3117 | 99.7 |
| 20 fps | $0.468+0.578t-0.0178t^2-0.0145t^3$ | 3200x2860 | 183 |

**Supplementary Figure 2: Automated calibration routine of voice coil actuator for higher speeds. a** FWHM vs. pixel rows graph used to determine uniformity of line profiles across a FOV. **b** Process of automated calibration of the voice coil for higher frame rates. **c** Table showing polynomial functions representing calibration curves for different frame rates and corresponding FOVs.

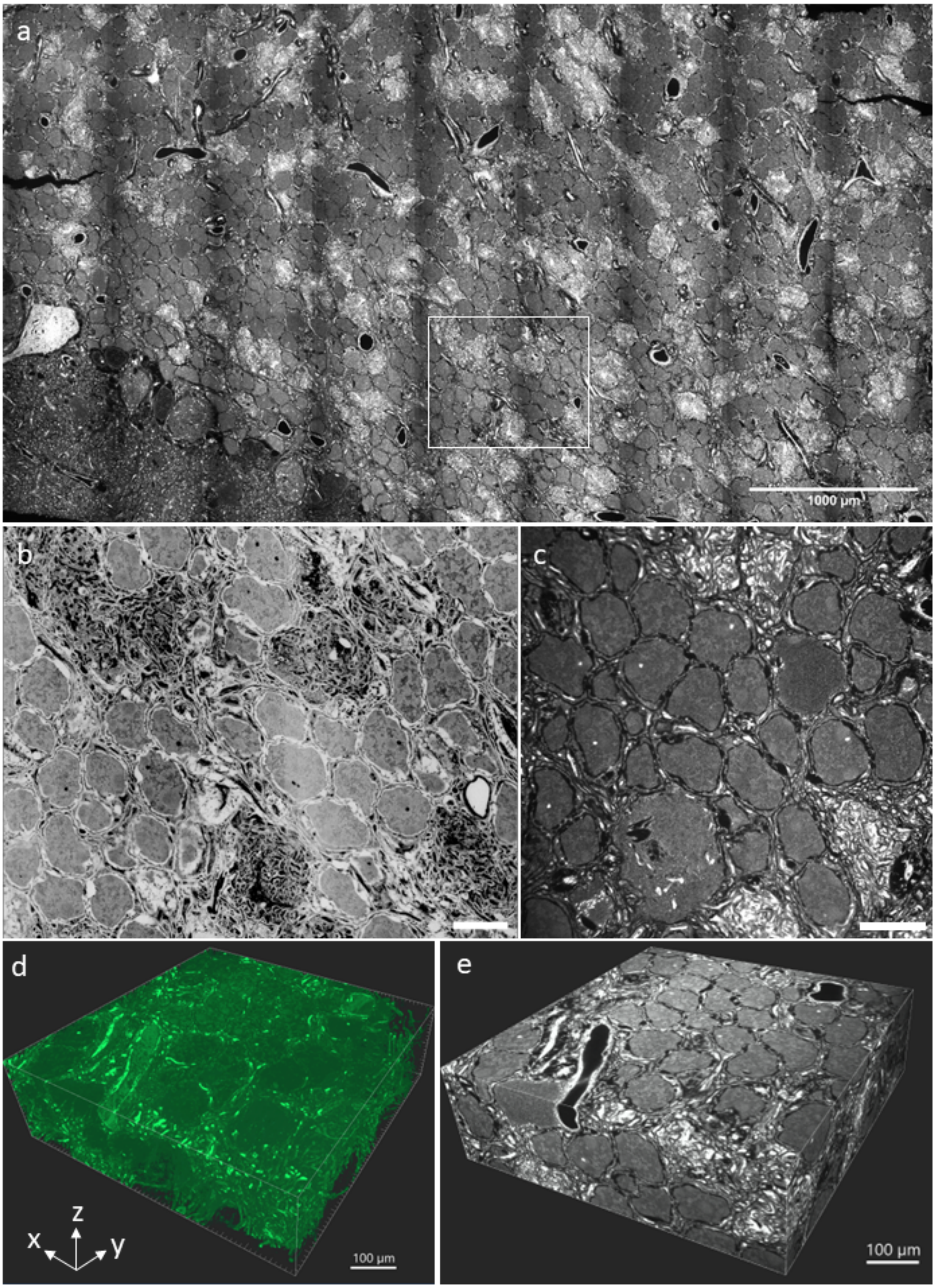


**Supplementary Figure 3: Large-FOV imaging of pan-expanded mouse brain tissue. a** Tiled pan-ASLM image of a 5.51 mm x 3.07 mm pan-expanded mouse brain tissue section stained with Atto 643 NHS ester. **b** Zoomed-in view of the region shown in (**a**) with inverted color table. **c** A single-FOV image showing pan-expanded brain tissue from the same dataset. **d** 3D rendering of brain tissue volume from the same dataset as (**c**). **e** 3D rendering of brain tissue volume from the same dataset as (**c**). Scale bars are not corrected for the expansion factor. Scale bars: (**b, c**) 100 µm.

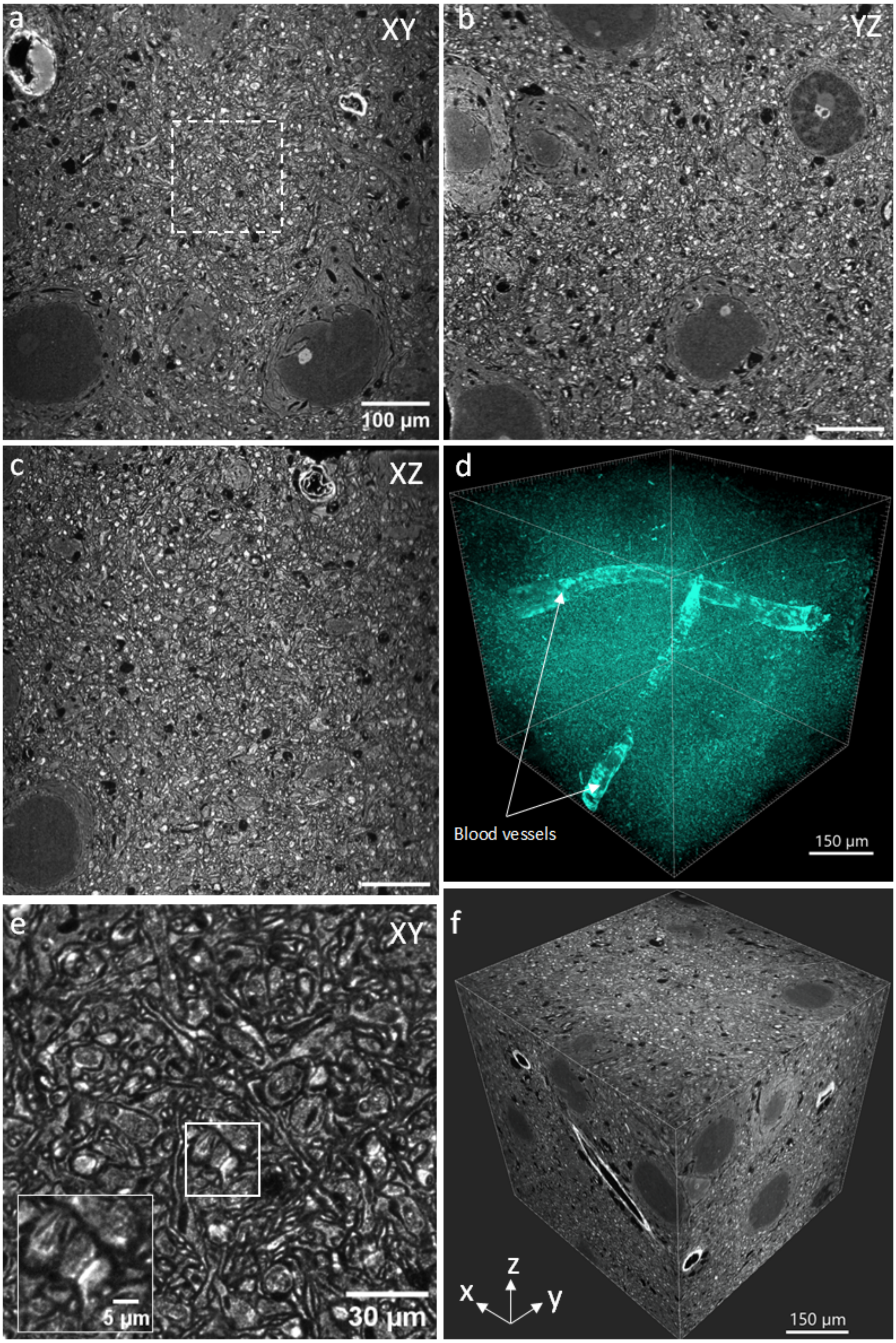


**Supplementary Figure 4: pan-ASLM imaging of pan-expanded mouse brain tissue. a** XY slice of a single-FOV image stack of a pan-expanded mouse brain tissue section stained with Atto 643 NHS ester. **b, c** YZ view (**b**) and XZ (**c**) of the same region as shown in (**a**). **d** MIP of the brain tissue volume shown in (**a**) revealing blood vessels. **e** Zoomed-in view of the region shown in the box in (**a**). The inset shows the synapse highlighted by the white box in (**e**). **f** 3D rendering of the brain tissue volume shown in (**d**). Scale bars are not corrected for the expansion factor. Scale bars: (**b,c**) 100 µm.

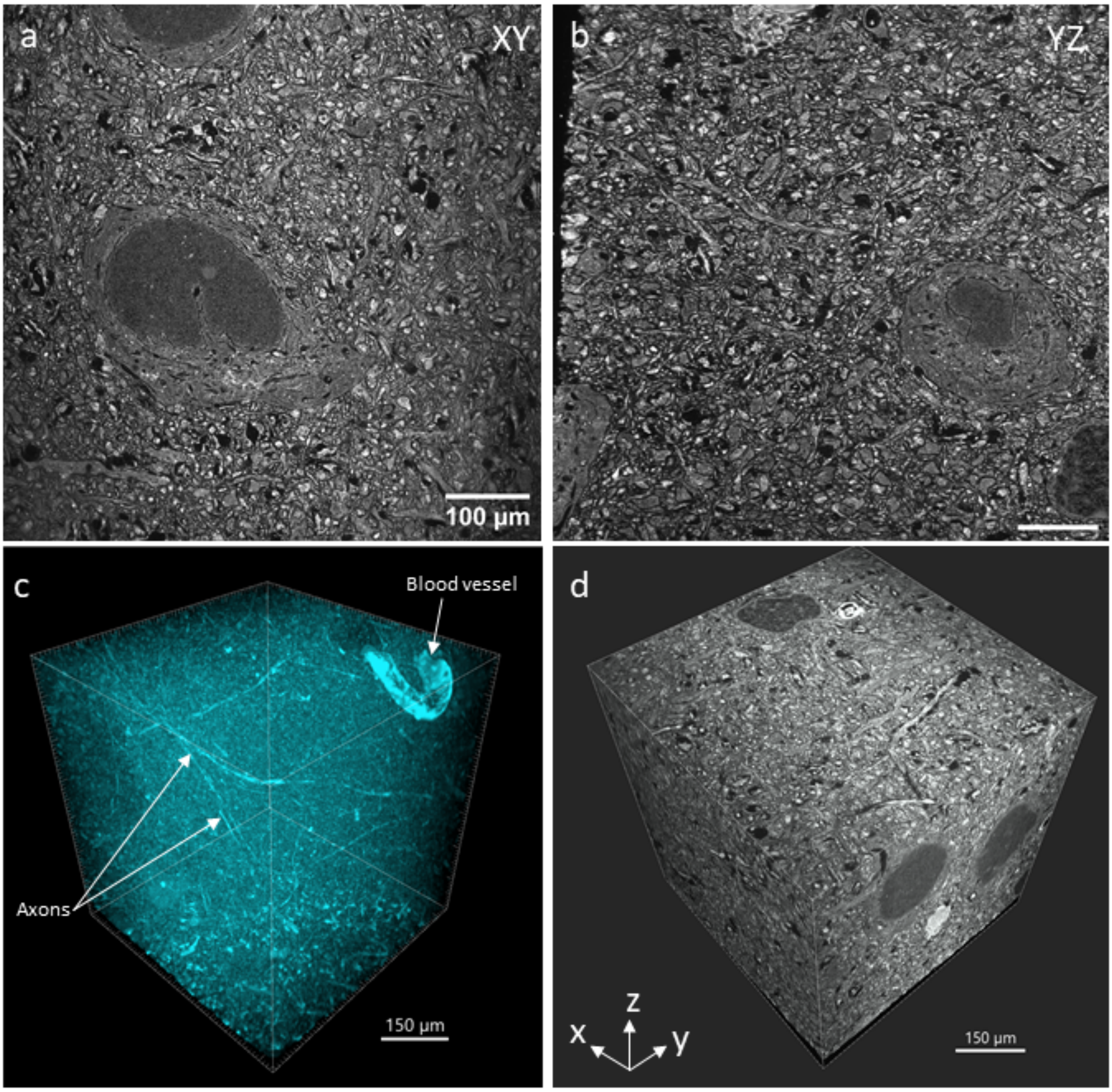


**Supplementary Figure 5: pan-ASLM imaging of pan-expanded mouse brain tissue. a** XY slice of a single-FOV image stack of a pan-expanded mouse brain tissue section stained with Atto 643 NHS ester. **b** YZ view of the same region as shown in (**a**). **c** MIP of the brain tissue volume shown in (**a**) revealing blood vessels and axons. **d** 3D rendering of the same brain tissue volume. Scale bars are not corrected for the expansion factor. Scale bar: (**b**) 100 µm.

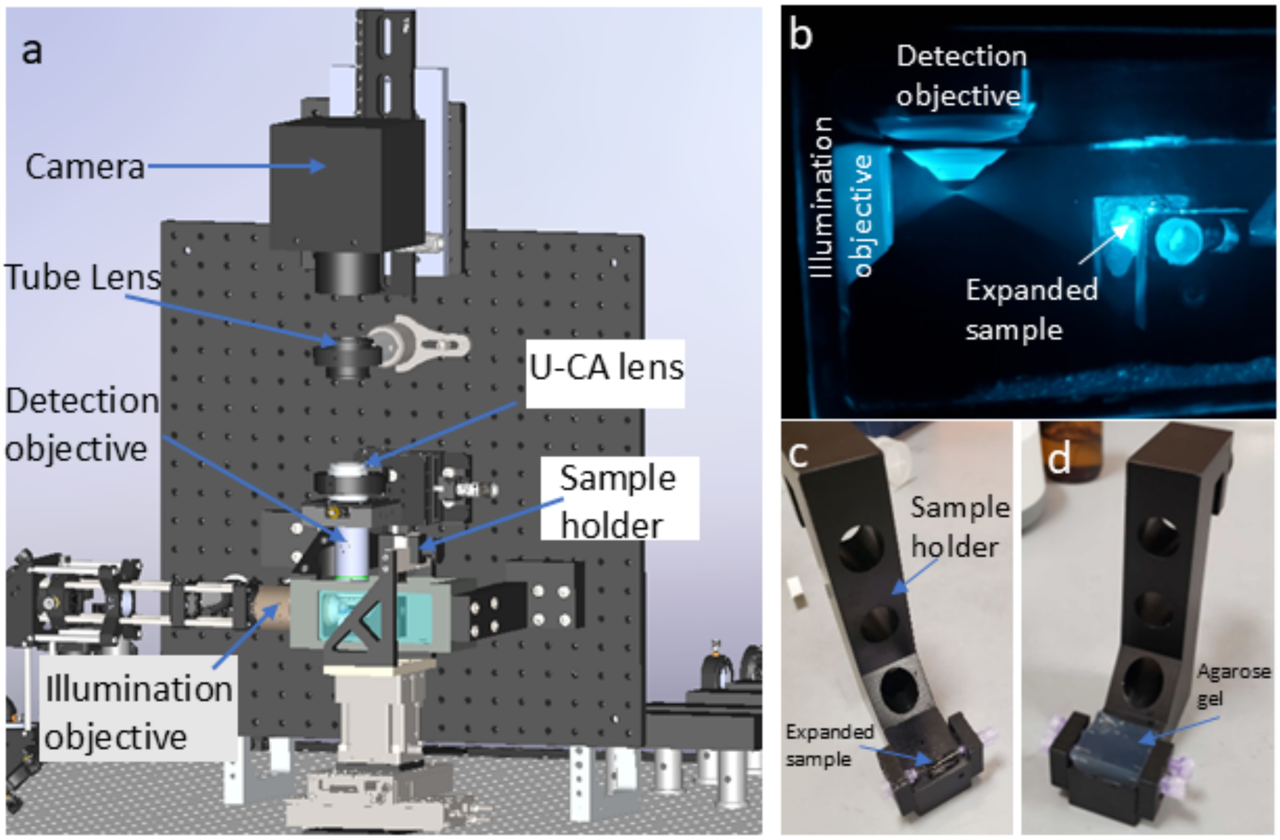


**Supplementary Figure 6: pan-ASLM microscope details. a** CAD design of the vertical breadboard of the pan-ASLM setup illustrating the detection beam path. **b** Image of the light sheet focused by the illumination objective before an imaging session. **c** Expanded sample mounted on the sample holder. **d** Agarose gel mounted on the sample holder.

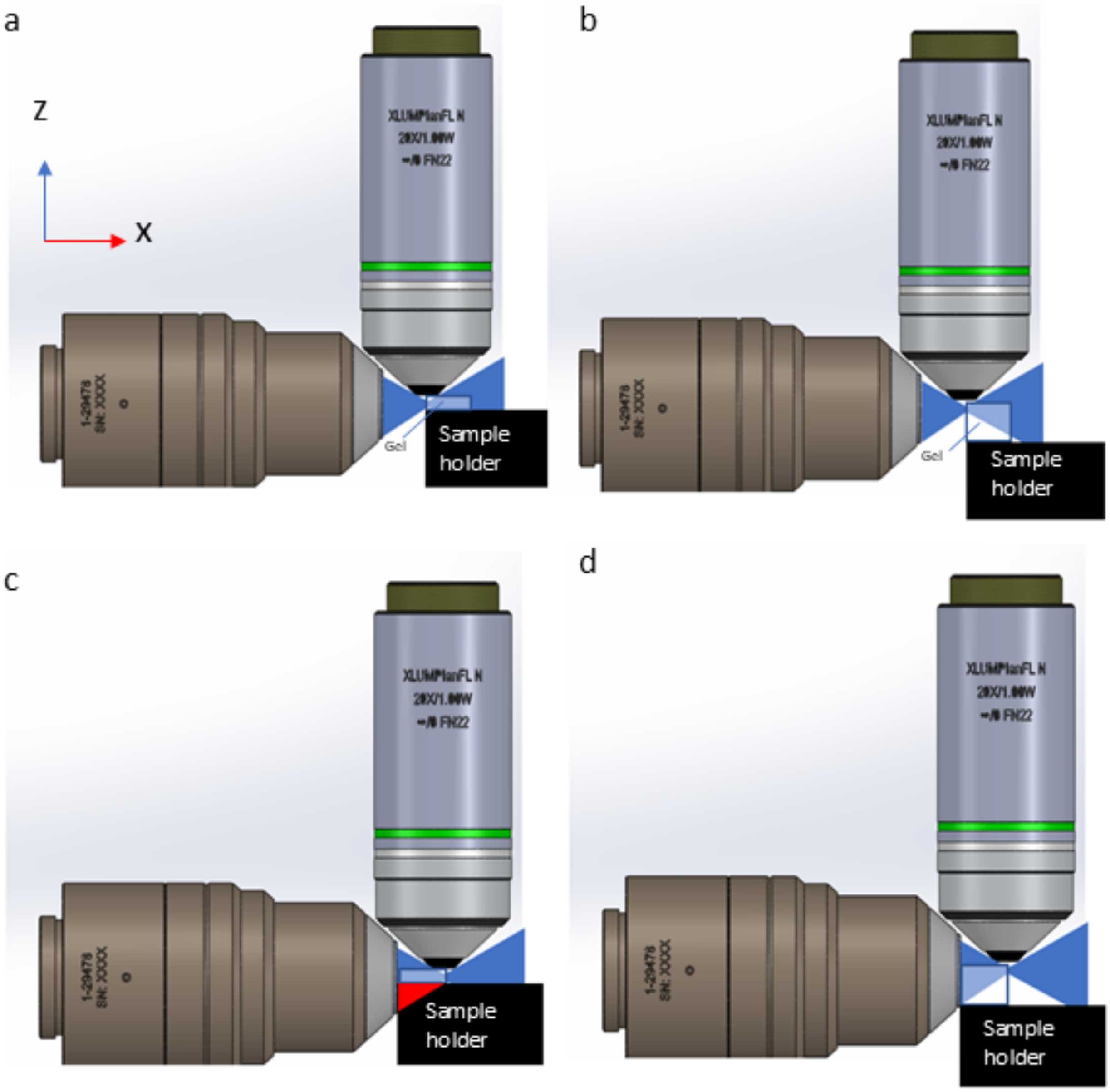


**Supplementary Figure 7: pan-ASLM imaging volume and sample thickness. a, b** The front of a thin (~2-mm thick) (**a**) and thick (~5 mm) sample (**b**) can be imaged with pan-ASLM across the whole depth. **c, d** When focusing deep into the sample with the illumination objective, the light sheet is clipped by the sample holder for the thin sample (**c**), but not the thick sample (**d**). Using a thick gel even for thin tissue samples or adding a spacer underneath a thin sample prevents this problem.

| Components | Company | Part number/ model name | Quantity |
|---|---|---|---|
| Laser(488 nm), 150mW | Coherent | OBIS LX 488nm 150mW | 1 |
| Laser(561 nm), 500mW | MPB Communications | 2RU-VFL-P-500-561-B1R | 1 |
| Laser(595 nm), 500mW | MPB Communications | 2RU-VFL-P-500-595-B1R | 1 |
| Laser(642 nm), 2W | MPB Communications | 2RU-VFL-P-2000-642-B1R | 1 |
| Dichroic mirror | Semrock | LM01-613-25 | 1 |
| Dichroic mirror | Semrock | Di02-R561-25x36 | 1 |
| Dichroic mirror | Semrock | Di02-R488-25x36 | 1 |
| AOTFnC-VIS | AA Optoelectronic | AOTFnC-VIS | 1 |
| AOTF driver, 8 channel | AA Optoelectronic | MPDS8C-B66-22-74-158 | 1 |
| Achromatic Half-Wave Plate | Thorlabs | AHWP05M-580 | 1 |
| f=50 mm Achromat | Thorlabs | AC254-50-A | 2 |
| f=300mm, Achromat | Thorlabs | AC254-300-A | 1 |
| piShaper | Edmund Optics | #12-644 | 1 |
| Periscope assembly | Thorlabs | RS99 | 1 |
| f=150mm, Achromat | Thorlabs | ACY254-150-A | 1 |
| f=100mm, Cylinderical lens | Thorlabs | ACY254-100-A | 1 |
| f=150mm, Achromat | Thorlabs | AC254-150-A | 3 |
| Polarizing beam splitter | Thorlabs | CCM1-PBS251/M | 1 |
| Quarter wave plate | Thorlabs | AQWP10M-580 | 1 |
| Remote focusing objective | Evident | UPLXAPO20X/0.8 NA | 1 |
| Scan mirror | Thorlabs | BB03-E02 | 1 |
| Voice coil actuator | Equipment Solutions | LFA-10 | 1 |
| Servo controller for voice coil | Servo controller | SCA814 | 1 |
| f=200mm, Achromat | Thorlabs | AC254-50-A | 1 |
| Illumination objective | ASI | 54-12-8 | 1 |
| Detection objective | Evident | XLUMPLFLN20XW | 1 |
| Piezo objective actuator | Thorlabs | PIA13 | 1 |
| Multiband emission filter | Chroma | ZET405/488/561/640mv2 | 1 |
| Magnification changer | Evident | U-CA | 1 |
| Tube lens | Evident | SWTLU-C | 1 |
| Camera | Teledyne Photometric | Kinetix | 1 |
| Vertical breadboard | Thorlabs | MBH4545/M | 1 |
| 18" Right-angle support | Thorlabs | VB01B/M | 2 |
| Coarse stage for sample (X,Y) | Physik Instrumente | V-508.231 | 2 |
| ACS controller | Physik Instrumente | G-901.R3197 | 1 |
| Translation Stage with Standard Micrometer | Thorlabs | MT1/M | 2 |
| Base Plate for MT Series Translation Stages | Thorlabs | MT401/M | 1 |
| Sample chamber | Tristar | Custom | 1 |
| Sample holder | Tristar | Custom | 1 |
| Stage baseplate | Tristar | Custom | 1 |
| 2nd stage baseplate | Tristar | Custom | 1 |
| 3rd stage baseplate | Tristar | Custom | 1 |

| | | | |
|---|---|---|---|
| Crossbar | Tristar | Custom | 1 |
| Sample chamber side mount | Tristar | Custom | 1 |
| Sample chamber upper mount | Tristar | Custom | 1 |
| Laser head mount | Tristar | Custom | 3 |
| AOTF mount | Tristar | Custom | 1 |

**Supplementary Table 1: pan-ASLM parts list.**

| Frame rate (fps) | Acquisition time (ms) | Flyback (ms) | Total time (ms) | 2 pixels exposure time (ms) | 4 pixels exposure time (ms) | 8 pixels exposure time (ms) |
|---|---|---|---|---|---|---|
| 1 | 976 | 24 | 1000 | 0.61 | 1.22 | 2.44 |
| 2 | 476 | 24 | 500 | 0.2975 | 0.595 | 1.19 |
| 2.5 | 376 | 24 | 400 | 0.235 | 0.47 | 0.94 |
| 5 | 176 | 24 | 200 | 0.11 | 0.22 | 0.44 |
| 10 | 76 | 24 | 100 | 0.0475 | 0.095 | 0.19 |
| 12.5 | 46 | 24 | 70 | 0.02875 | 0.0575 | 0.115 |
| 20 | 26 | 24 | 50 | 0.01625 | 0.0325 | 0.065 |

**Supplementary Table 2: Table showing exposure times corresponding to each scan width and frame rate.**
